# Testing for local adaptation and evolutionary potential along 1 altitudinal gradients in rainforest *Drosophila*: beyond laboratory estimates

**DOI:** 10.1101/068080

**Authors:** Eleanor K. O’Brien, Megan Higgie, Alan Reynolds, Ary A. Hoffmann, Jon R. Bridle

## Abstract

Predicting how species will respond to the rapid climatic changes predicted this century is an urgent task. Species Distribution Models (SDMs) use the current relationship between environmental variation and species’ abundances to predict the effect of future environmental change on their distributions. However, two common assumptions of SDMs are likely to be violated in many cases: (1) that the relationship of environment with abundance or fitness is constant throughout a species’ range and will remain so in future, and (2) that abiotic factors (e.g. temperature, humidity) determine species’ distributions. We test these assumptions by relating field abundance of the rainforest fruit fly *Drosophila birchii* to ecological change across gradients that include its low and high altitudinal limits. We then test how such ecological variation affects the fitness of 35 *D. birchii* families transplanted in 591 cages to sites along two altitudinal gradients, to determine whether genetic variation in fitness responses could facilitate future adaptation to environmental change. Overall, field abundance was highest at cooler, high altitude sites, and declined towards warmer, low altitude sites. By contrast, cage fitness (productivity) *increased* towards warmer, lower altitude sites, suggesting that biotic interactions (absent from cages) drive ecological limits at warmer margins. In addition, the relationship between environmental variation and abundance varied significantly among gradients, indicating divergence in ecological niche across the species’ range. However, there was no evidence for local adaptation within gradients, despite greater productivity of high altitude than low altitude populations when families were reared under laboratory conditions. Families also responded similarly to transplantation along gradients, providing no evidence for fitness trade-offs that would favour local adaptation. These findings highlight the importance of (1) measuring genetic variation of key traits under ecologically relevant conditions, and (2) considering the effect of biotic interactions when predicting species’ responses to environmental change.

## INTRODUCTION

Understanding the factors that determine species’ distributions and local abundances is a central goal of ecology, and is essential for predicting how populations, species and ecological communities will respond to environmental change (Ehrlén & Morris, 2015). Species’ distribution models (also known as ecological niche or bioclimatic envelope models) are used to relate species’ abundances to environmental variables, and to predict shifts in their distributions based on future climatic conditions (Elith & Leathwick, 2009, Guisan & Thuiller, 2005, Pearson & Dawson, 2003, Thomas *et al.*, 2004). Such models typically assume that the association between the environment and a species’ abundance (i.e. its niche) does not vary across the species’ geographical range, and will remain stable in the future (but see Kearney *et al.*, 2009). However, spatial variation in environmental tolerances is observed across many species’ ranges, demonstrating local niche differentiation (Banta *et al.*, 2012, Kelly *et al.*, 2012). In addition, genetic variation within populations may generate rapid evolutionary responses to environmental change *in situ*, allowing population persistence beyond current ecological limits (Bridle & Vines, 2007, Hoffmann *et al.*, 2015, Hoffmann & Sgrò, 2011).

Ignoring variation in a species’ ecological niche within populations, or between populations across its geographical range, will have two contrasting consequences: (1) we may overestimate the geographical distribution of a species if tolerances are assumed to be constant throughout the species’ range (i.e. that all populations can tolerate all currently occupied conditions: Hampe, 2004, Kelly *et al.*, 2012); and (2) we may underestimate the potential for species to persist through evolutionary change, where extinction would be predicted based on current distributions (Davis *et al.*, 2005, Hoffmann & Sgrò, 2011, Kearney *et al.*, 2009). Understanding the potential for rapid adaptation generated by standing genetic variation in fitness, both among and within populations, is therefore crucial when predicting the impacts of environmental change on population persistence, and the future geographical distributions of species (Atkins & Travis, 2010, Chevin *et al.*, 2010, Hampe, 2004, Holt, 2009, Lavergne *et al.*, 2010).

Studies testing for local adaptation and genetic variation in environmental tolerances in the context of predicting responses to environmental change are rare for animals, where attention has focused on the evolution of traits in single populations (e.g. Charmantier & Gienapp, 2014, Kruuk *et al.*, 2008). These data are more widely available in plants, and have been used to project future responses to environmental change. For example, Banta *et al.* (2012) modelled the niche breadth of *Arabidopsis thaliana* genotypes that varied in flowering time, and found a more than four-fold difference between genotypes in the size of their potential distributions. Similarly, studies of local adaptation in forest trees reveal genetic divergence in phenology and other ecological traits that are associated with their broad geographical distributions (e.g. Alberto *et al.*, 2013, Kremer *et al.*, 2012). In the few cases where genetic variation in ecological traits has been estimated across multiple populations in animals, this has typically been done under controlled conditions in the laboratory, rather than under field conditions, which will vary far more in time and space, meaning that selection may act on many more traits simultaneously, or at different points in time. Because environmental conditions affect the heritability of many traits (Charmantier & Garant, 2005, Hoffmann & Merilä, 1999, Kruuk *et al.*, 2008), laboratory assays of genetic variance in traits or fitness may not predict evolutionary trajectories in natural populations (Pemberton, 2010). These issues mean there is an urgent need for data on genetic variation in fitness across a range of naturally varying environments, to determine how the relationship between the environment and fitness varies due to local adaptation, or in relation to genetic variation within populations.

*Drosophila birchii* is endemic to the tropical rainforests of north-eastern Australia and Papua New Guinea (Schiffer & McEvey, 2006). Laboratory assays of environmental tolerance traits in this species have revealed genetic divergence along both latitudinal (Griffiths *et al.*, 2005, Hoffmann *et al.*, 2003, van Heerwaarden *et al.*, 2009) and some altitudinal (Bridle *et al.*, 2009) gradients, consistent with local adaptation to temperature and humidity variation. In addition, laboratory assays have revealed lower levels of genetic variation in ecologically important traits associated with tolerance of climatic stresses within populations close to the species’ range margin, which may constrain adaptation (e.g. Hoffmann *et al.*, 2003, Kellermann *et al.*, 2006). These results suggest that ecological tolerances vary substantially throughout the range of *D. birchii*, and that the potential for adaptation to environmental change also varies among populations. However, genetic variation in fitness under field conditions has not previously been measured, therefore it is not known how predictions of evolutionary potential based on genetic variation in traits measured in the laboratory relate to fitness variation in the more variable field environment, where biotic interactions are common and complex, and are themselves mediated by variation in abiotic factors.

In this study, we examine the relationship between local abundance of *D. birchii* and environmental variation along four altitudinal gradients. These altitudinal gradients represent local ecological limits of this species, and show temperature and humidity variation across distances of 4-16 km of a similar magnitude to that observed across hundreds of kilometres of latitudinal range (see Table S1). In addition, we transplanted families of laboratory-reared *D. birchii* in cages along two altitudinal gradients and tracked their fitness under naturally-varying environmental regimes, in order to: (i) determine the effect of environmental change (simulated by movement along an environmental gradient) on fitness of *D. birchii*, (ii) test for local adaptation across these gradients, and (iii) estimate genetic variation in fitness, both overall, and in response to movement along the gradient (i.e., genetic variation in the ‘reaction norms’ of fitness). By transplanting virgin flies, we ensured that courtship, mating, reproduction, and the development and survival of offspring occurred entirely under field conditions, and therefore captured all of these important components of fitness variation. Flies in cages experienced abiotic conditions similar to those outside cages, but were not exposed to biotic interactions. Therefore, by comparing the change in fitness of *D. birchii* in cages as a result of movement along environmental gradients with the change in its field abundance, we were also able to test the degree to which abiotic environmental conditions alone determine species’ distributions. Furthermore, by transplanting flies from multiple populations and families, we were able to evaluate the role of among-population divergence in mediating this relationship, and the potential for rapid changes in ecological tolerances in the future through adaptation. Finally, by comparing laboratory estimates of genetic variation in fitness with those made in the field, we provide one of the first tests of how trait genetic variation estimated in the laboratory predicts the potential for evolutionary responses to environmental change under more ecologically realistic conditions.

## MATERIALS AND METHODS

### *Predicting the local abundance of* D. birchii *from environmental variables*

#### *Estimating* D. birchii *abundance along altitudinal gradients*

Adult *D. birchii* were collected between February-May in 2010-2012 from a total of 94 sites, comprising 10-30 sites along each of four altitudinal gradients (Paluma, Kirrama, Mt Edith and Mt Lewis) in northern Queensland, Australia. Gradients were between 16°30’S and 19°00’S latitude (a distance of ~300 km), and spanned altitudes from 23–1233m above sea level (a.s.l.), over distances of 3.7 –16.3 km (Figure 1; Table S1). At each site, 5–20 buckets of mashed banana (> 1 day old) were placed at least 5m apart for 5–10 days. Flies were sampled from each bucket twice daily using a sweep net; captured flies were then sorted under CO_2_ anaesthesia to identify *D. birchii,* and to isolate *D. birchii* females for isofemale line generation (see below).

**Figure 1.**
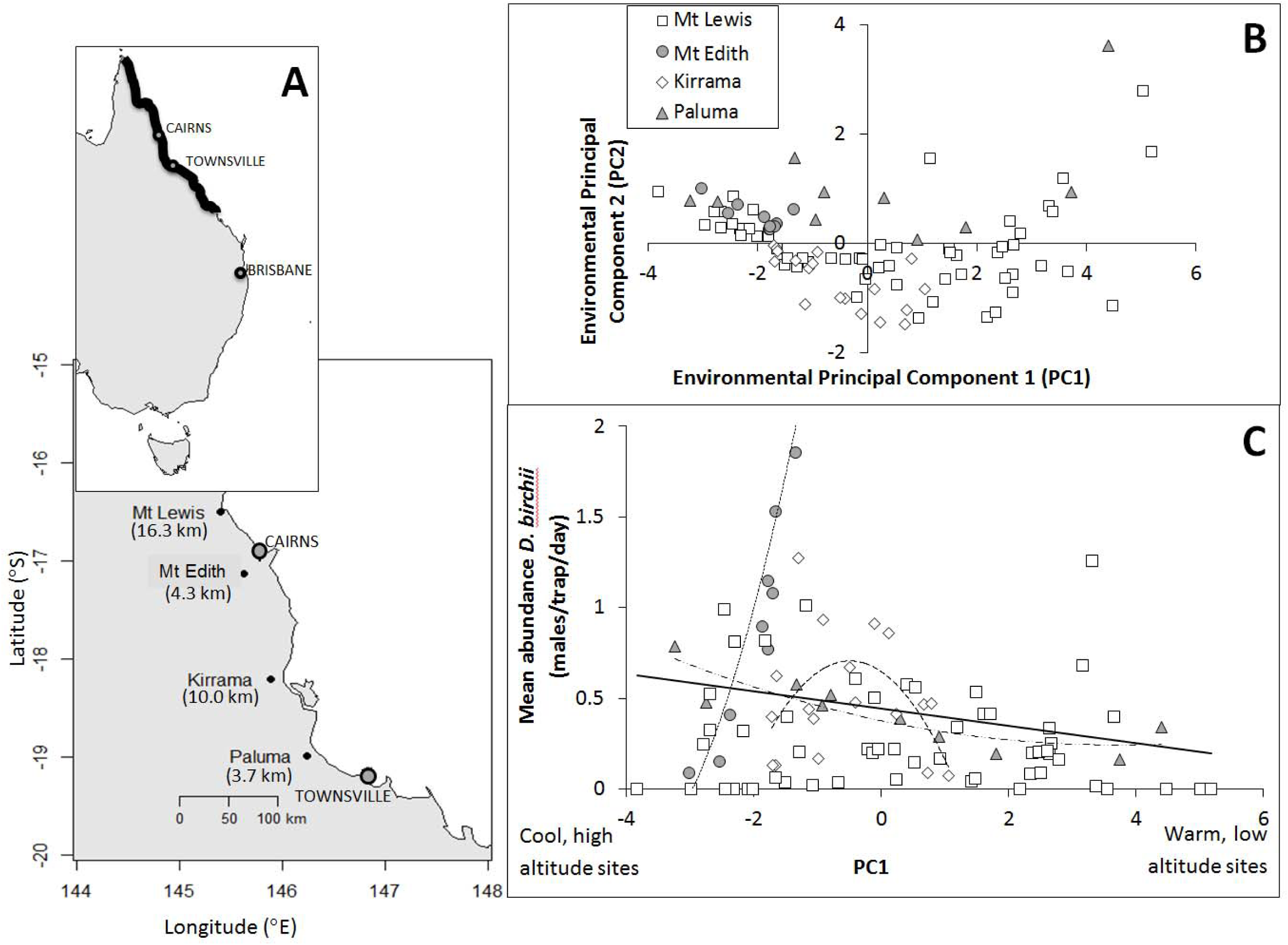
**A.**Locations of the altitudinal gradients where *D. birchii* was sampled, with the length of each gradient given in brackets. Thick black line on top map indicates the extent of *D. birchii*’s distribution in Australia. **B.** Plot showing the position of sites along each of the four altitudinal gradients with respect to the first two Principal Components (PC1 and PC2) from a Principal Components Analysis of six environmental variables. The loading of each environmental variable on these PCs is shown in Table S4. **C.** Mean abundance of *D. birchii* against PC1 for each of the four altitudinal gradients (see legend in B for interpretation of symbols). Fitted curves from linear models of mean abundance on PC1 are shown for gradients where this relationship was significant (*P*< 0.05; see Table 1): Mt Edith (dotted line), Kirrama (dashed line) and Paluma (dash-dot line). The solid line is the fitted relationship from a model including the full data set (*R*^2^ = 0.10, *F*_1,90_= 10.68, *P*= 0.002).

Estimates of local abundance were the mean number of *D. birchii* males captured per site per day, as used by Bridle *et al.*(2009). We used the number of males captured (rather than total number of flies) because female *D. birchii* cannot be distinguished from closely-related species in the *serrata* species complex, *D. serrata* and *D. bunnanda*, whereas males can be identified by examining their genital bristles (Schiffer & McEvey, 2006). Females can only be identified by examining their male offspring, therefore using females in abundance counts would bias estimates towards those females that successfully reproduced. We estimated abundance at 48 sites along two gradients sampled in 2010 (Kirrama and Mt Lewis) and 46 sites along three gradients in 2011 (Paluma, Mt Edith and Mt Lewis) (Figure 1; Table S1). There was no significant variation in the magnitude or distribution of abundance between the two years of sampling at Mt Lewis (the only gradient sampled in both years; see Table S1), therefore abundance data at sites along this gradient were combined across years.

#### *Measuring environmental predictors of* D. birchii *abundance*

Tinytag dataloggers were attached to trees 1.5 – 1.8 m above the ground at 10 – 30 sites along each altitudinal gradient to take hourly measurements of temperature (°C) and % relative humidity (RH) between February 2010 and June 2012. This included the sampling period, as well as the duration of the cage transplant experiments (see below). In addition, the abundance of the other *serrata* complex species, *D. bunnanda* and *D. serrata*, was estimated at each site based on numbers of males captured in traps. This variable was included to provide a measure of the frequency of interspecific interactions at different points along gradients. These species are closely related to *D. birchii*, use similar resources, and have partially overlapping geographical distributions, although their local abundances show different patterns with respect to environmental conditions (Schiffer & McEvey, 2006). *Drosophila serrata* has a much broader latitudinal range than *D. birchii*, and is considered a habitat generalist. *Drosophila bunnanda*, like *D. birchii*, is a rainforest specialist and has a more restricted distribution, with a southern border more than 500 km north of that of *D. birchii*. Neither of these species was present at Mt Edith, but they were found at some sites at Paluma, Kirrama and Mt Lewis. At the sites sampled, *D. bunnanda* was much more common than *D. serrata* (determined by genotyping field-caught males at the diagnostic locus *Eip 75B*), and numbers of *D. serrata* captured were too low to be used as an independent predictor of *D. birchii* abundance. We therefore combined estimates of the abundances of *D. bunnanda* and *D. serrata* as a single measure.

Temperature and humidity data from Tinytag dataloggers and estimates of the abundance of other species of the *serrata* species complex were used to produce six environmental predictors of *D. birchii* abundance: (a) abundance of non-*D. birchii serrata*-complex species (NONBIRCH), (b) mean daily minimum temperature (MDT_min_), (c) mean daily temperature (MDT), (d) mean daily maximum temperature (MDT_max_), (e) mean daily temperature range (MDT_diff_), and (f) mean daily relative humidity (MDH).

Linear regression revealed that most of the six environmental variables were strongly correlated with both latitude and altitude (Table S2). The environmental variables were also all highly correlated with one another (Table S3). To avoid collinearity of factors in models predicting abundance, we identified a set of uncorrelated variables by conducting a Principal Components Analysis (PCA) using the *prcomp* function in *R* v3.1.2 (R Core Team, 2014), with all variables standardised to a mean of 0 and standard deviation of 1. Temperature and humidity data collected over the full two-year measurement period were used in the PCA. These values were highly correlated with those seen over the sampling periods, and over the course of the caged transplant experiments. The first two Principal Components (PCs) together accounted for 89% of the total variation in the environmental variables (Table S4), and were included as factors in linear models predicting *D. birchii* abundance. The first Principal Component (PC1) captured the majority (76.8%) of variation in the environmental variables, with relatively even loadings of all six variables, whereas PC2 was dominated by the abundance of other *serrata*-complex species (Table S4). The positions of sites at each gradient with respect to PC1 and PC2 are shown in Figure 1.

A linear model was fitted using the full set of abundance data across all gradients, with mean *D. birchii* abundance at each site as the response variable, and the following terms included as predictors: gradient (categorical variable with four levels corresponding to the four altitudinal gradients), linear and polynomial (quadratic) terms for PC1 and PC2 (continuous variables), and interactions of gradient with each of the linear and quadratic PC terms. Abundance data were weighted by the number of sampling days at each site in the linear model. We fitted an additional set of models for each gradient separately, to explore environmental predictors found to differ among gradients in their relationship with *D. birchii* abundance in the full model. Linear models were fitted using the *lm* function in *R* v3.1.2 (R Core Team, 2014).

### Testing for genetic variation in responses to environmental change: cage transplant experiment

In March-May 2012, 35 isofemale lines from two sites at the top and bottom of both the Mt Edith and Paluma gradients were collected, and reared through two generations to large numbers under laboratory conditions. They were then subjected to a line-cross design within collecting sites (see below and Figure S1), and virgin males and females from the lines generated were transplanted into 591 cages at multiple sites along the altitudinal gradient from which they were originally sampled (Figure S2). Total productivity was assessed for each cage, allowing tests for local adaptation and estimates of genetic variation in fitness at each gradient under naturally-varying environmental conditions. Because virgin flies were placed in cages *in situ* at field sites, all courtship, mating, egg-laying and larval and pupal development occurred under naturally varying conditions. Despite being of similar length, the steepness and altitudinal ranges of the gradients differ. Paluma is much steeper than Mt Edith, covering twice the altitudinal range, and a much broader range of temperatures, humidity, and abundance of *serrata*-complex species (Table S1), as captured by the first two PCs (Figure 1). The design of the experiment is illustrated in Figure S2; details of the experimental procedures are given below.

#### Establishment of isofemale lines

Individual field-mated females collected from two high and two low altitude sites at Paluma and Mt Edith were placed in 40 ml glass vials with 10 ml standard *Drosophila*media (agar,raw sugar,inactive yeast,propionic acid and methyl-4-hydroxybenzoate), supplemented with live yeast, and left to oviposit for four days to initiate isofemale lines. These mothers were transferred to a fresh food vial every four days until they no longer produced offspring. Offspring of the same mother were then mixed across vials to found the next generation. The genital bristles of the male offspring of each female were examined to distinguish *D. birchii* from the morphologically similar sympatric species *D. bunnanda* and *D. serrata*. Five *D. birchii* isofemale lines were established for each site (four sites per gradient; 20 lines in total per gradient), and each isofemale line was maintained across 2–4 vials in a constant temperature (CT) room at 25 °C on a 12:12 hour light:dark cycle.

#### Breeding flies for cage transplant experiment

Isofemale lines collected from field sites were maintained in the laboratory for two generations after collection from the field in order to standardise maternal environment effects. Following this, we established crosses between lines from the same site to ensure rapid generation of large numbers for field transplants (Figure S1). We paired virgin females from each line with virgin males from each of the other lines from the same site (i.e. excluding within-line crosses), with three replicates per line-cross combination. The crossing scheme used to generate flies for cage transplants is summarised in Figure S1.

Each pair was left to mate and lay for five days. Offspring emerging from these crosses were counted and sexed on eclosion (± 12 hrs) each day until emergence was complete, and flies held separately by sex (up to 10 flies per vial) for up to 10 days before being transplanted to field cages. We then pooled offspring from the same maternal isofemale line, keeping the sexes separate to ensure all flies were unmated prior to establishing cages. Flies transplanted into cages therefore ranged in age from 3 – 10 days, but mixing together flies from the same maternal isofemale line meant that their distribution across cages and sites was random with respect to age. We used this approach to avoid excluding lines with low fecundity from being tested in the field. Transplanting ‘maternal isofemale lines’ (hereafter referred to as ‘lines’) rather than generating mass bred lines for each site allowed us to maintain representation in our experiment of as many maternal lines as possible, as well as (crucially) enabling partitioning of among-line (genetic) variation in fitness under field conditions.

#### Establishment of field transplant cages

The cages used for field assays of line fitness were 600 ml clear plastic bottles with two 135 mm × 95 mm windows cut out, covered with 2 mm fly wire mesh and 30-denier nylon stocking material, which allowed movement of air through the cages. Each cage was encased in 20 mm wire mesh to prevent attack by birds and mammals. This cage construction allowed the survival and productivity of flies to be monitored, while exposing them to temperature and humidity that were as close to naturally-varying conditions as possible. We dispensed 90 ml of media (as described above) directly into the bottom of each cage. This volume of food was 9 times that used to rear offspring of the same number of flies at low density in the laboratory (see methods of line maintenance above), to prevent food becoming a limiting resource during this experiment, and to minimize density-dependent competition among larvae. Cages were suspended from tree branches between 1.5–1.8 m above the ground. We placed iButton temperature loggers (Maxim integrated Products, San Jose, CA, USA) inside five of the cages at each site to record temperature hourly, to test that temperatures within cages were consistent with those measured outside by the Tinytag data loggers, and to assay temperature variation among cages within sites. The iButtons revealed low variability in temperature within, relative to between sites (90% of variation in mean temperature was between compared to within sites), and Tinytag and cage temperature measurements were highly correlated (*R^2^*= 0.88, *P* <0.001). Figure S3 shows a comparison between iButton measurements inside cages and Tinytag measurements outside cages for mean daily temperature (MDT), mean daily minimum temperature (MDT_min_) and mean daily maximum temperature (MDT_max_). Linear models comparing the two measures revealed no significant difference between measures inside and outside of cages for MDT or MDT_max_, although measurements of MDT_min_were, overall, lower inside cages than outside at field sites (Figure S3). It is likely that the positioning of cages (hung from tree branches), compared with that of data loggers (attached to tree trunks) meant that cages were slightly more exposed, resulting in lower minimum temperatures inside cages. There was no significant difference between the two measures for the change in MDT, MDT_min_ or MDT_max_in relation to altitude along gradients (Figure S3). The iButtons did not measure relative humidity (RH), therefore it was not possible to compare RH inside and outside cages. While it is likely that RH in cages was increased relative to the outside air, mean daily RH was high at all sites (RH > 74%, and usually RH >88%; Table S1), therefore we consider that RH is unlikely to be a limiting factor for survival and reproduction of *D. birchii*.

Lines were transplanted only to sites along their gradient of origin, not between gradients. At each gradient, cage locations included the two high and two low altitude sites from which the lines were collected, as well as sites at intermediate altitudes (Figure S2). At Mt Edith, 15 lines (9 from low altitude, 6 from high altitude sites) were transplanted along the gradient. At Paluma, 20 lines (10 from each end of the gradient) were transplanted. However, due to variation in fecundity of lines in the laboratory, there were insufficient flies to transplant all lines to all sites at each gradient. At Mt Edith, between 9 – 15 lines were transplanted at each site, and this always included both high and low altitude lines (Figure S2). At Paluma low altitude lines had much lower fecundity than high altitude lines (see below and Figure S4). To maximise power to detect local adaptation (see below), Paluma lines from both high and low altitude populations were transplanted to cages at the two high and two low altitude sites from which lines were sourced (18 – 19 lines per site; Figure S2), but only high altitude lines were transplanted to intermediate sites (6 – 8 lines per site; Figure S2). We established 325 cages at nine sites at Mt Edith (mean = 36.1 cages/site), and 266 cages at ten sites at Paluma (mean = 26.6 cages/site) (Figure S2). Five virgin male and female flies from the same line were placed in a given cage. At each site there were 2 – 4 cages per line. Exact numbers of lines and cages transplanted to each site along each gradient are shown in the table within Figure S2.

#### Estimates of fitness of flies in cages

We monitored each cage daily for five days after establishment and recorded the number of surviving adult flies each day. On the fifth day, we removed all surviving flies to ensure they were not included in offspring counts used to measure productivity (see below). We then left cages *in situ* for another 25 days (30 days total) to allow offspring to pupate and hatch, even at the coolest sites. After 30 days, all cages were taken to the laboratory, where they were held for five days at 25°C to ensure that all offspring had emerged from that generation. The first offspring did not emerge until after 20 days at any site, while the majority of offspring had emerged at all sites by day 30, therefore the emerging offspring were all from a single generation. Total productivity (number of offspring emerging) was used as a measure of fitness for each cage. This includes the effects of parental survival, however mean survival was high (Mt Edith = 75.2%; Paluma = 80.8%) and did not vary significantly along either gradient, or among lines, therefore the majority of productivity variation was driven by variation in reproductive success. The short lifespan and relatively low population density of *D. birchii* means that mating opportunities are likely to be a major factor limiting the lifetime fitness of *D. birchii.* This, combined with the high and uniform survival of flies in cages along altitudinal gradients, means that early fertility is likely to be a very important component of fitness variation in this species. Therefore, while further data would be required to evaluate fitness variation at later life history stages, we argue that within the logistical constraints of such a large experiment, focusing on this measure of fitness is justified.

#### Analysis of fitness variation in field cages

We fitted generalised linear mixed models (GLMMs) analysing variation in fitness (productivity) in cages along each gradient to: (1) test for local adaptation, and (2) estimate genetic variation in fitness, and in the effect of movement along a gradient on fitness (‘reaction norms’ in fitness of lines), in order to estimate the potential for adaptive responses to environmental change.

To test for local adaptation, we used the ‘sympatric-allopatric’ (SA) contrast proposed by Blanquart *et al.* (2013). This method compares the fitness of *sympatric* populations (populations transplanted back to their site of origin) with that of *allopatric* populations (populations transplanted to a different site from their site of origin), while controlling for variation due to habitat (i.e., environmental variation among transplant sites) and source population (i.e., due to genetic differences in fitness among source populations) (Blanquart *et al.*, 2013). This comparison has greater power to detect local adaptation than other more restrictive definitions of local adaptation (e.g., the ‘home vs away’ and ‘local vs foreign’ comparisons described by Kawecki and Ebert (2004)) (Blanquart *et al.*, 2013). Power to detect local adaptation using this method increases as a function of the number of sympatric-allopatric comparisons, which for a given number of transplants is maximised by transplanting all source populations back into the source sites. We additionally tested for variation in fitness reaction norms along gradients, which required transplanting lines to a larger number of sites (including sites that had not been used as source populations). Nevertheless, by ensuring that *D. birchii* from all lines within a gradient were transplanted to gradient ends (where flies were sourced), we still had high power to detect local adaptation within the constraints imposed by these dual aims of our experiment.

GLMMs included as fixed effects: (1) environmental variables (a subset of PC1, PC2, (PC1)^2^, and (PC2)^2^. Terms were sequentially removed and models compared to determine whether each improved model fit; see results), to account for habitat variation among transplant sites, (2) ‘source population’, a categorical variable with four levels corresponding to the populations from which *D. birchii* were sourced within a gradient, and (3) a ‘local adaptation’ term indicating whether a cage was ‘sympatric’ or ‘allopatric’, as defined above. Evidence for local adaptation is indicated by significantly higher fitness of sympatric cages than allopatric cages, after controlling for habitat and population effects.

We included random intercept and slope terms for the effect of line (nested within source population) to estimate: (i) genetic variation in fitness (averaged across the whole gradient), and (ii) variation among lines in fitness responses to environmental change (“fitness reaction norms”), respectively. Random slope terms tested for variation in the fitness responses of lines with respect to the same environmental variables as were included as fixed effects in the model (i.e. a subset of PC1, PC2, (PC1)^2^, and (PC2)^2^; see above and results).

Productivity data were over-dispersed relative to the Poisson distribution generally used for modelling count data, and had an excess of zeroes due to over-representation of cages from which no offspring emerged. We therefore modelled productivity as a negative binomial distribution (Lindén & Mãntyniemi, 2011), specifying zero-inflation, and used a log link function. GLMMs were fitted using the *R* package *glmmADMB* 0.8.0 (Fournier *et al.*, 2012, Skaug *et al.*, 2013). Separate models were fitted for each gradient.

#### Genetic variation in productivity in the laboratory

We assessed variation among lines and source populations from Paluma and Mt Edith in their productivity in the laboratory for comparison with genetic variation estimated from field cages. Productivity was measured as the number of offspring emerging from crosses established to generate flies for the caged transplant experiment (see above), therefore it included the same set of lines as in analyses of fitness variation in field cages. We again fitted GLMMs using *glmmADMB*, with the same distribution as in analyses of fitness variation in cages (see above). We included source population as a fixed predictor, and maternal isofemale line (nested within source population) as a random factor. To assess whether lines with high productivity under laboratory conditions also performed well in the field, we compared the rank order of lines for productivity in the laboratory and in the field using a Spearman’s rank correlation test, implemented using the *cor.test* function in *R* v3.1.2 (R Core Team, 2014). Separate models were fitted for each gradient in both sets of analyses.

### *Predicting local abundance of* D. birchii *from variation in fitness in cages*

We fitted linear models to test how well fitness in cages predicted local abundance of *D. birchii* at the gradients where caged transplants were undertaken (Paluma and Mt Edith). We used mean productivity in field cages as a measure of fitness at each site. Fitness and abundance data were both standardised to a mean of 0 and standard deviation of 1 so that they were on the same scale. We fitted linear models with the *lm* function in *R* v3.1.2 (R Core Team, 2014), using standardised productivity as the predictor variable, and standardised abundance of *D. birchii* as the response variable. Separate models were fitted for each gradient.

## RESULTS

### *Predicting local abundance of* D. birchii *from environmental variables*

At all gradients except Mt Lewis, the first Principal Component (PC1) from the PCA of environmental variables was a significant predictor of *D. birchii* abundance (Table 1; Figure 1C). However, the strength and shape of the relationship between PC1 and abundance varied substantially between gradients (Table 1; Figure 1C). Abundance of *D. birchii* increased with PC1 at Mt Edith (indicating increased abundance at higher temperatures/lower altitudes), and decreased with PC1 at Paluma (Table 1; Figure 1C). At Mt Edith and Kirrama, model fit was improved by the addition of a quadratic term for PC1 (Table 1). Given that the four gradients span different altitude and temperature ranges (Table S1), these different patterns reflect, in part, variation in the range of values of PC1 present within each gradient (Figure 1B). However, differences are still evident when gradients are compared over equivalent values of PC1 (Figure 1C). PC2 did not improve the fit of the model of *D. birchii* abundance overall (Table 1), or of models of *D. birchii* abundance within each gradient.

**Table 1.** Predicting *D. birchii* abundance along four altitudinal gradients based on environmental variation. Environmental variation is represented by the first two principal components (PC1 and PC2) of the ordination analysis (see text and Figure 1B). For the overall analysis, gradient and interactions of gradient with linear and quadratic terms for each predictor were included. Factors showing a significant interaction with gradient (PC1 and PC1^2^) were then included in models for each gradient individually. Model statistics indicating the fit of each model are also shown. Significant terms (p < 0.05) in each model are in *italics*.

| Predictor | df | SS | F | P | Model statistics |
| --- | --- | --- | --- | --- | --- |
| Gradient | 3 | 20.57 | 9.99 | $1.39 \times 10^{-5}$ | Adj. $R^2 = 0.404$ |
| PC1 | 1 | 2.10 | 3.06 | 0.09 | $F_{19,72} = 4.25$ |
| PC1<sup>2</sup> | 1 | 0.85 | 1.23 | 0.27 | $P = 3.86 \times 10^{-6}$ |
| PC2 | 1 | 0.73 | 1.06 | 0.31 |  |
| PC2<sup>2</sup> | 1 | 0.25 | 0.36 | 0.55 |  |
| Gradient x PC1 | 3 | 17.78 | 8.64 | $5.70 \times 10^{-5}$ | |
| Gradient x PC1<sup>2</sup> | 3 | 8.82 | 4.29 | 0.01 |  |
| Gradient x PC2 | 3 | 3.14 | 1.53 | 0.22 |  |
| Gradient x PC2<sup>2</sup> | 3 | 1.19 | 0.58 | 0.63 |  |
| Residual | 72 | 49.42 |  |  |  |

***Estimates of parameters for each gradient:***
| Gradient | Parameter | Estimate (SE) | t | P | Model statistics |
| --- | --- | --- | --- | --- | --- |
| Mt Lewis | Intercept | 0.264 (0.046) | 5.722 | $5.28 \times 10^{-7}$ | Adj. $R^2 = 0$ |
| | PC1 | -0.015 (0.021) | -0.721 | 0.474 | $F_{2,52} = 0.892$ |
| | PC1<sup>2</sup> | -0.003 (0.007) | -0.430 | 0.669 | $P = 0.416$ |
| Mt Edith | Intercept | 6.094 (1.144) | 5.328 | 0.002 | Adj. $R^2 = 0.920$ |
| | PC1 | 4.052 (1.057) | 3.833 | 0.009 | $F_{2,6} = 46.82$ |
| | PC1<sup>2</sup> | 0.683 (0.234) | 2.923 | 0.027 | $P = 0.0002$ |
| Kirrama | Intercept | 0.650 (0.108) | 6.043 | $2.25 \times 10^{-5}$ | Adj. $R^2 = 0.213$ |
| | PC1 | -0.247 (0.109) | -2.254 | 0.040 | $F_{2,15} = 3.295$ |
| | PC1<sup>2</sup> | -0.254 (0.102) | -2.490 | 0.025 | $P = 0.065$ |
| Paluma | Intercept | 0.373 (0.046) | 8.185 | $7.88 \times 10^{-5}$ | Adj. $R^2 = 0.681$ |
| | PC1 | -0.071 (0.016) | -4.517 | 0.003 | $F_{2,7} = 10.58$ |
| | PC1<sup>2</sup> | 0.009 (0.006) | 1.513 | 0.174 | $P = 0.008$ |

### Testing for genetic variation in responses to environmental c 1 hange: caged transplant experiment

#### Testing for local adaptation along altitudinal gradients

There was no evidence for local adaptation within gradients; ‘sympatric’ cages did not outperform ‘allopatric’ cages after controlling for habitat and population effects at either gradient (Table 2; Figure 2). At Mt Edith, the SA contrast was only marginally non-significant (*P*=0.052; Table 2), but fitness of allopatric cages exceeded that of sympatric cages (Figure 2), which is opposite to expectations if the difference is due to local adaptation. At Paluma, there was no significant difference between the fitness of sympatric and allopatric cages, and the trend was also opposite to that predicted with local adaptation (*P*=0.774; Table 2; Figure 2).

**Figure 2.**
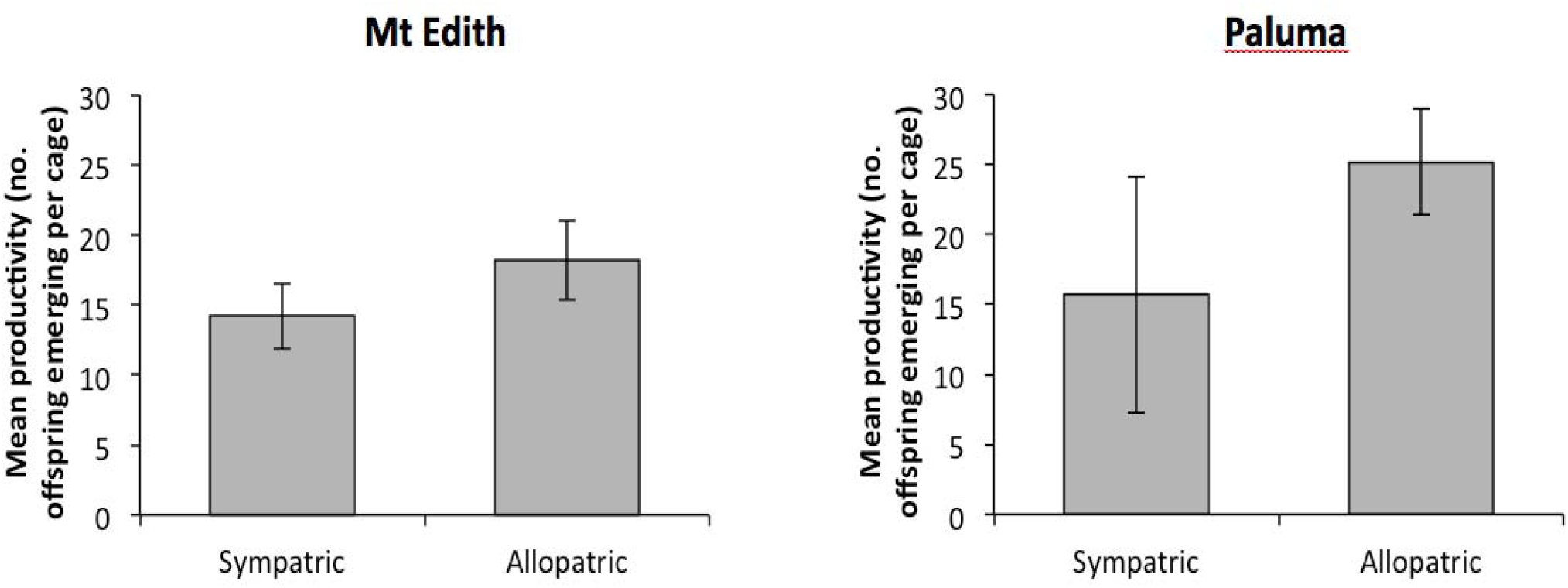
No evidence for local adaptation in caged transplants. Plots show the results of tests for local adaptation in caged transplants at Mt Edith and Paluma using sympatric-allopatric (SA) contrasts. The mean productivity (no. offspring emerging) of cages of flies transplanted back into their site of origin (sympatric), and those transplanted to all of the other sites along the same gradient (allopatric) are shown. Error bars are standard errors across the four source populations when transplanted sympatrically or allopatrically. The difference in productivity between sympatric and allopatric populations was marginally non-significant at Mt Edith (*P*= 0.052) and non-significant at Paluma (*P*= 0.774) (see Table 2).

**Table 2.**
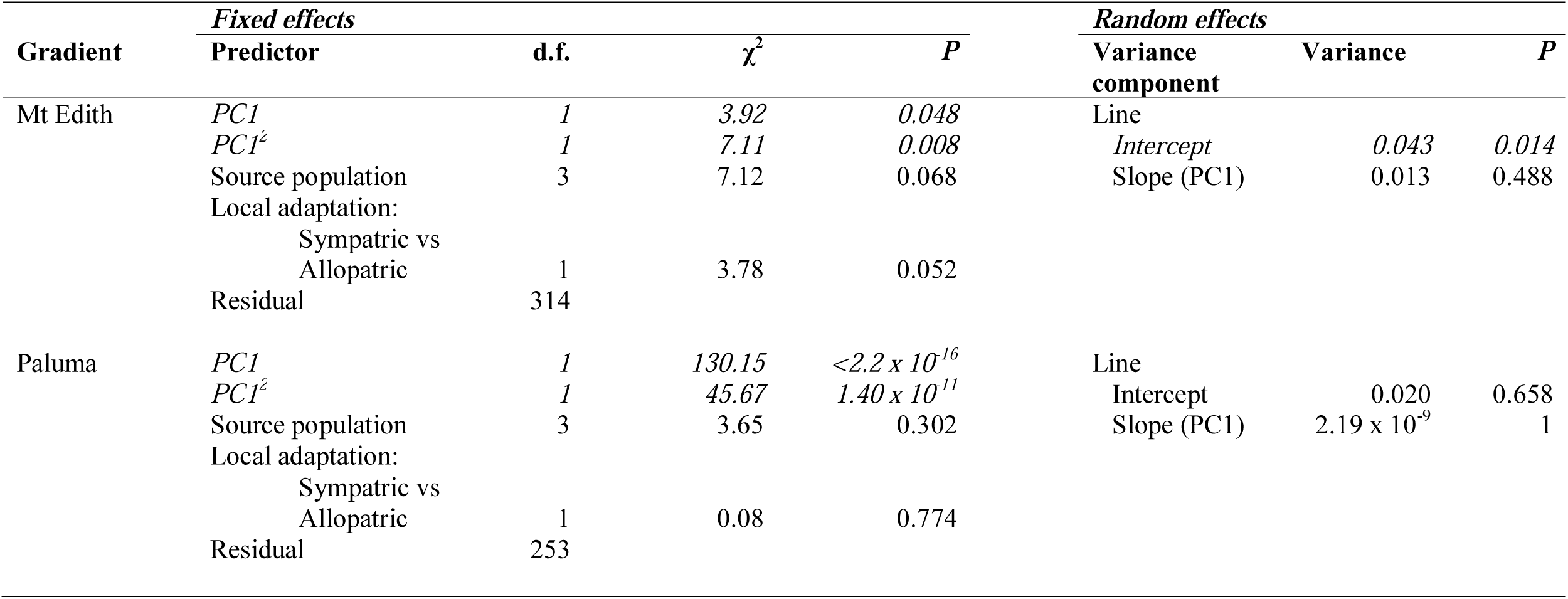
Tests for local adaptation and genetic variation in fitness from caged transplant experiments along the Mt Edith and Paluma altitudinal gradients, using Generalised Linear Mixed Models (GLMMs). The fixed effects included: linear and quadratic terms for PC1, which were significant predictors of *D. birchii* abundance at these gradients, ‘source population’, and a ‘local adaptation’ term which compared cages transplanted back to the site where flies originated (‘sympatric’) with those transplanted to a different site (‘allopatric’). Random intercept and slope (with respect to PC1) terms for the effect of isofemale line nested in source population (‘Line’ in table) were also included. Significant effects are denoted in *italics*. The significance of fixed effects was evaluated using a χ^2^ test, and of random effects using a likelihood-ratio test comparing models with and without each term included. Variance components were estimated after removing non-significant fixed effects from the model.

There were highly significant effects of environmental variation on fitness in cages. Along both altitudinal gradients there was a significant, non-linear increase in cage productivity with increasing PC1 (increasing temperature) (Figure 3). Source population effects approached significance at Mt Edith (*P*=0.068; Table 2), which was attributable to low fitness of flies from one of the source populations (Figure S5), and was non-significant at Paluma (*P*=0.302; Table 2; Figure S5).

**Figure 3.**
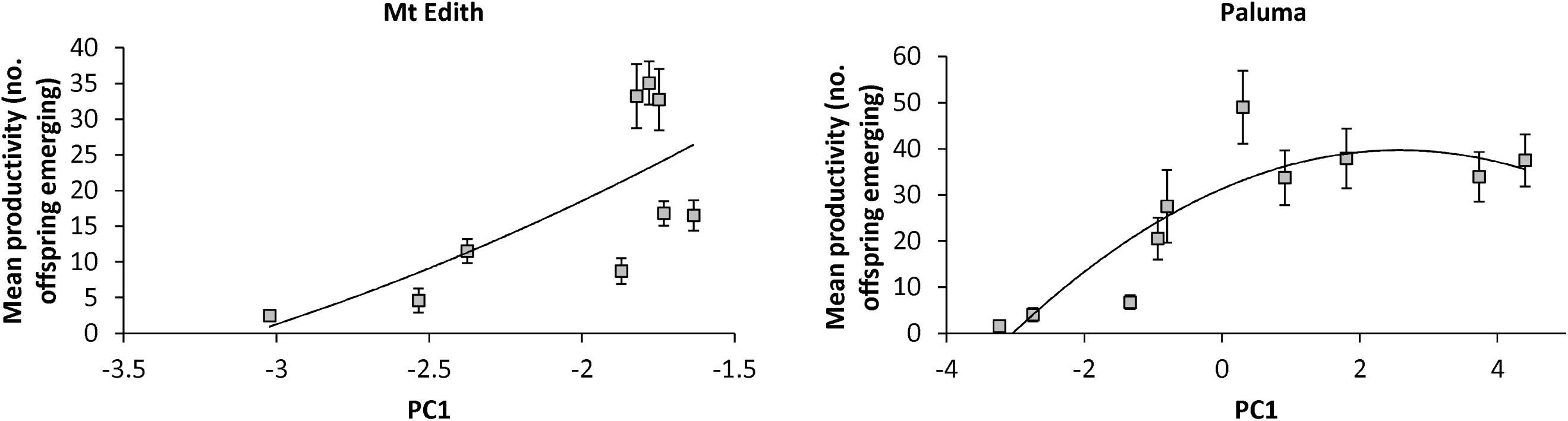
Fitness in cages increases with mean temperature along altitudinal gradients. Plots show mean productivity of cages placed at each e along altitudinal gradients at Mt Edith and Paluma. Mean site productivity (averaged across the cages at each site) is plotted as a function of PC1, the first principal component of a PCA of variation for a set of environmental variables (see Methods and Figure 1B), which is strongly, positively associated with temperature. Fitted curves are from linear models of productivity on PC1 for each gradient (see Table 2). Error bars indicate standard errors based on isofemale lines at each site. Note that the Paluma gradient encompasses a much wider range of values of PC1 than Mt Edith. Points have been offset slightly along the x-axis at Mt Edith to reduce overlap.

#### Variation in fitness and reaction norms of fitness among lines

There was significant variation among lines in their productivity in cages at Mt Edith (*P* = 0.014), but not at Paluma (*P* = 0.658)(Table 2). At Mt Edith, the mean productivity of the ‘fittest’ line (24.5 offspring/cage) was more than seven times that of the least fit line (3.4 offspring/cage), whereas at Paluma the fittest line (37.5 offspring/cage) had mean productivity twice that of the least fit (19 offspring/cage). We did not detect significant variation among lines in the slopes of their responses (i.e. their ‘reaction norms’ of fitness) to the change in environment experienced as a result of being transplanted along gradients, as captured by variation in the slopes of their fitness with respect to PC1 (Table 2). Random slope variation with respect to the other PC terms was also not significant for either gradient. These results suggest that there is significant genetic variation in mean fitness across these environmental conditions at Mt Edith, but at both gradients all lines respond similarly to the change in environment; that is, lines with high relative fitness at one end of the gradient tend to have high fitness at all sites.

#### Genetic variation in productivity in the laboratory

Consistent with results from the field experiment, we found significant among-line variation in laboratory productivity at Mt Edith, but not Paluma (Table S5). Estimates of among-line variance in the laboratory were higher than in the field for both gradients (Table S5; *cf* Table 2), although the crossing scheme may have reduced genetic and maternal effect differences between the lines in field cages. In contrast to the field experiment, variation among source populations for laboratory productivity was highly significant at both gradients (Table S5), with high altitude source populations showing higher productivity than low altitude populations in both cases (Figure S4). A Spearman’s rank correlation test revealed that, while the rank order of lines for productivity in the laboratory and in the field was positively correlated at both gradients, the correlation was not distinguishable from zero at either gradient (Mt Edith: ρ = 0.271, *P* = 0.327; Paluma: ρ = 0.173, *P* = 0.492), suggesting the relative fitness of lines under constant conditions is not a good predictor of their relative fitness in the more variable field environment. However, note again that productivity in the field was measured on flies from line crosses, as opposed to measurements on isofemale lines in the laboratory.

### *Predicting local abundance of* D. birchii *from fitness in cages*

Productivity of *D. birchii* in field cages changed in the same direction as local abundance of *D. birchii* along the Mt Edith gradient, and this relationship was marginally non-significant (Slope (SE) = 1.313 (0.57); *P* = 0.054). However, this relationship was significantly negative along the Paluma gradient (Slope (SE) = −0.253 (0.08); *P* = 0.012) (Table S6; Figure 4). As outlined above, productivity in cages increased with increasing PC1 (i.e., towards warmer, lower altitude sites) at both gradients (Figure 3). Paluma covered a wider range of PC1 values than Mt Edith; specifically, Paluma included much higher values, reflecting higher temperatures. Therefore, the difference between the gradients in the relationship between fitness in cages and abundance implies that, while cage productivity is a good predictor of local abundance of *D. birchii* at cooler, high altitude sites, it fails to predict changes in abundance towards the warm margin of this species’ range.

**Figure 4.**
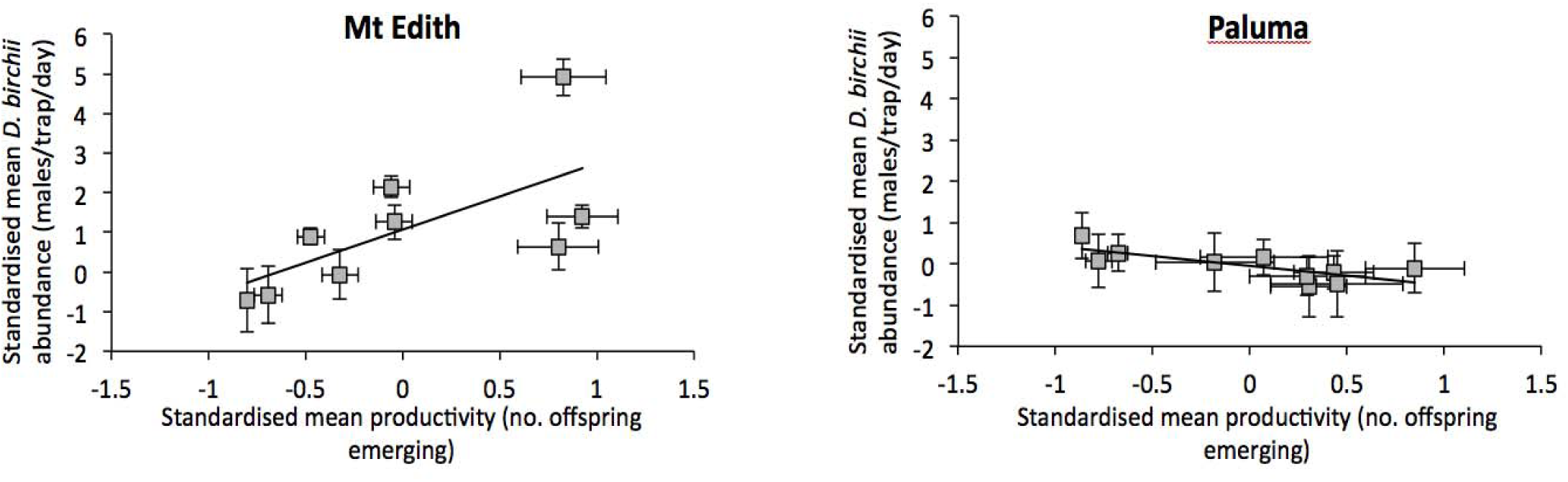
Cage fitness predicts local abundance at cool, high altitude sites but not at warm, low altitude sites. Plots show the relationship tween fitness estimated from the caged transplant experiment (cage productivity) and the local abundance of *D. birchii* estimated from field mpling at Mt Edith and Paluma. Fitness and abundance data were both standardised to mean = 0 and standard deviation = 1. Error bars on undance (y-axis) are standard errors across sampling days, and on productivity (x-axis) are standard errors among lines. Fitted lines are shown m regressions of standardised mean *D. birchii* abundance on standardised mean productivity (see Table S4).

## DISCUSSION

Predicting the effect of rapid environmental change on species’ distributions, and therefore the persistence of ecological communities, is an urgent priority. However, such predictions typically rely on models that assume a constant relationship between abiotic environmental variation and species’ persistence or abundance, thus ignoring the potential for evolutionary change in environmental tolerances, and the influence of biotic interactions. Our approach, which combines surveys of field abundance, cage transplant experiments, and both laboratory and field estimates of genetic variation in fitness in the rainforest fruit fly *Drosophila birchii*, provides a comprehensive test of these assumptions, along ecological gradients that characterise distributional limits of this species at different spatial scales.

### *Predicting responses to environmental change from the relationship between* D. birchii *abundance and environmental variation*

Our field surveys revealed that local abundance of *D. birchii* is strongly predicted by environmental variation at three of the four altitudinal gradients studied, which each exhibits variation in mean temperature characteristic of hundreds of kilometres of latitudinal distance (Table S1). Overall, there was a decline in the abundance of *D. birchii* towards warm, low altitude sites (Figure 1), which suggests that the rising temperatures forecast as a result of climate change will reduce the area of suitable habitat for this species. However, the relationship between environment and local *D. birchii* abundance differed between gradients (Table 1), suggesting local variation in the response of this species to environmental change, at least across the period measured here. Predictions of *D. birchii* abundance based on its association with environmental variables at a broad geographical scale may therefore perform poorly at a local scale. This variation in the relationship of *D. birchii* abundance with environmental conditions could be caused by other factors affecting abundance that vary among gradients that were not captured by our measures of environmental variation, and/or local adaptation within or among gradients, enabling population growth over different ranges of environments at different gradients. We consider each of these possibilities below.

### *Cage transplants along altitudinal gradients: does the abiotic environment predict the fitness of* D. birchii*?*

Fitness, as measured by productivity in cages, showed consistent increases with temperature along both gradients. This was in contrast to the reduction in abundance at warmer (low altitude) sites in our surveys of field abundance. This surprising result suggests that there are factors excluded from our cages that restrict *D. birchii*’s distribution at its warm ecological limit. The cage transplant experiment exposed flies to changes in the naturally-varying abiotic (i.e. temperature and humidity) environment, but there are likely to be significant changes in the biotic environment (e.g., competitors, predators, parasites, pathogens) over this scale that were absent from cages, and which may constrain *D. birchii*’s abundance towards its warmer margin. This is consistent with the hypothesis, initially proposed by Darwin (1859), and subsequently supported by numerous authors (e.g. Ettinger *et al.*, 2011, MacArthur, 1972), that abiotic factors are the principal limit to species’ distributions at high latitudes and altitudes, while the importance of biotic interactions increases towards warmer margins at lower latitudes and altitudes. The lowest latitude, and on average warmest, gradient included in our abundance survey, Mt Lewis, was the only gradient where abiotic environmental variation (captured by PC1) did not predict *D. birchii* abundance (Table 1), again suggesting a potential role for biotic factors. Further work is underway to identify important biotic interactions. However, we note that PC2, which is largely driven by the abundance of non-*birchii serrata*-complex species (Figure 1B), did not predict *D. birchii* abundance at any gradient (Table 1), suggesting that competition with these closely related species is not the key factor limiting the distribution of this species.

Understanding how biotic and abiotic factors interact to shape species’ distributions is crucial for predicting the responses of ecological communities to environmental change (Alexander *et al.*, 2015, Araújo & Luoto, 2007, Godsoe *et al.*, 2015, Grassein *et al.*, 2014). Predicting the effect of changes in either the abiotic or biotic environment on species distributions is complicated by the fact that these different components of environmental variation are typically highly correlated in nature. Most species’ distribution models either ignore biotic variables, or implicitly assume that these correlations will remain constant in future (Araújo & Luoto, 2007). However, abiotic and biotic factors may become uncoupled if interacting species within an ecological community differ in their responses to environmental change, resulting in novel species’ assemblages (e.g. Alexander *et al.*, 2015). Future studies should explicitly test for the effects of biotic interactions within and among species on fitness, in combination with abiotic factors, to better understand local variation in evolutionary responses to environmental change, and therefore the persistence of species and local communities in response to ongoing climate change.

### Local adaptation and genetic variation in fitness and reaction norms in response to movement along altitudinal gradients, and comparison with laboratory estimates

We did not detect evidence of local adaptation within either gradient within the timeframe of our caged transplant experiments. Although there was significant genetic variation in overall fitness at Mt Edith, all lines transplanted at both gradients responded similarly to the imposed change in their environment. In other words, reaction norms for fitness of different lines do not intersect or vary in steepness, indicating that fitness under conditions at one end of the gradient does not ‘trade off’ against fitness at the opposite end. The lack of local adaptation within gradients is surprising, because divergent selection between gradient ends is expected to be strong in this system, given the substantial and consistent difference in their abiotic environments (temperature and humidity), and the significant consequences of this for fitness of *D. birchii,* as shown by our cage transplant experiments. Possible explanations for a lack of local adaptation along gradients include gene flow, which has been shown to be high in this species over larger geographic distances than were considered here (Schiffer *et al.*, 2007, van Heerwaarden *et al.*, 2009), and may swamp local adaptation, particularly given the steep changes in abundance observed even between adjacent sites, which are likely to lead to asymmetrical gene flow (Bridle *et al.*, 2009, Bridle & Vines, 2007). Alternatively, populations occupying marginal habitat towards the species’ range edge may lack sufficient genetic variation to track local optima by adaptation, potentially due to small population size, or trade-offs between different components of fitness (Blows & Hoffmann, 2005). Differences in the relative importance of abiotic and biotic factors at each end of the altitudinal range of *D. birchii* may also explain why we did not detect either genetic variation in fitness reaction norms or local adaptation in our cage transplant experiment. If biotic interactions (rather than temperature or humidity) constrain the distribution of *D. birchii* at its warm margin, fitness trade-offs may become apparent when measured in the presence of such interactions. Finally, we note that fitness along the gradient was only measured on one occasion, whereas selection pressures can change across years ((Kingsolver *et al.*, 2001)) and should ideally be characterised repeatedly.

Nevertheless, previous studies in *D. birchii* have revealed latitudinal clines (over similar temperature ranges) suggestive of local adaptation in development time (Griffiths *et al.*, 2005), resistance to desiccation (Hoffmann *et al.*, 2003, Kellermann *et al.*, 2006) and starvation (Griffiths *et al.*, 2005, van Heerwaarden *et al.*, 2009), as well as altitudinal clines in chill coma tolerance (Bridle *et al.*, 2009). However, all of these studies examined trait variation under constant conditions in the laboratory. While it is likely that the patterns of trait variation they observed were the result of selection, the fitness consequences of this variation may become evident only under certain sets of conditions, since environmental conditions are known to affect estimates of trait heritabilities (Charmantier & Garant, 2005, Hoffmann & Merilä, 1999, Pemberton, 2010). We also found significant genetic variation in productivity among *D. birchii* populations in the laboratory, but not in the field. Importantly, the mean productivity of *D. birchii* in field cages was substantially lower than productivity in the laboratory, confirming a common assumption that laboratory conditions are benign relative to the conditions experienced by wild populations. A consequence of this may be that genetic differences among populations are not realised under less favourable field conditions due to masking by environmental variation. This highlights the importance of assaying fitness under naturally-varying conditions when inferring adaptive potential in wild populations. Furthermore, the timing and location of such studies should encompass conditions that are *a priori* thought to be most limiting for the focal species, to ensure that key drivers of selection are included.

### Implications for predicting biological responses to environmental change

Three important findings emerge from our study that enable evaluation of the accuracy of predicted changes in the distribution of *D. birchii* in response to environmental change using traditional Species’ Distribution Models. (1) The relationship between environmental variation and abundance differs between gradients, demonstrating the importance of geographic scale in predictive models. (2) The effect of abiotic environmental variation on fitness of *D. birchii* in cages does not mirror the change in field abundance, suggesting an important role for biotic interactions in limiting the distribution of this species. (3) There is no local adaptation nor genetic variation in fitness reaction norms of *D. birchii* within gradients, although this contradicts predictions based on laboratory estimates of genetic variation in fitness. These observations are likely to have general significance beyond the model system examined here, and can therefore offer insights on how to improve methods for predicting biological responses to environmental change.

Incorporating spatial geographic scale into Species’ Distribution Models is quite straightforward, as long as abundance or occurrence data are available at a sufficiently fine scale. Ideally, sampling should be undertaken across both local and global ecological limits, to account for potential variation in the factors limiting species’ distributions at these different scales (e.g. across altitudinal and latitudinal gradients Halbritter *et al.*, 2013). As has been appreciated by others, biotic interactions should be incorporated into SDMs by including data on the presence or abundance of co-occurring species as predictive factors (Araújo & Luoto, 2007, Wisz *et al.*, 2013). Our results demonstrate that the importance of biotic interactions in limiting species’ distributions is likely to vary across abiotic gradients, which reiterates the importance of sampling at appropriate geographic scales. Furthermore, given that key biotic interactions are themselves susceptible to the effects of changes in the abiotic environment, regular re-sampling should be undertaken to identify changes in the correlation of abiotic and biotic components, and their consequences for species’ distributions.

The lack of genetic variation in fitness reaction norms suggests that populations of *D. birchii* along gradients are likely to respond similarly to a changing thermal environment, and have low potential for adaptation. This contrasts with measurements under laboratory conditions (both in the present study and in previous work e.g. Bridle *et al.*, 2009), which reveal significant genetic variation in ecologically important traits both within and among populations sampled from different parts of the species’ altitudinal range. These data highlight the importance of assessing genetic variation in fitness under ecologically relevant conditions when predicting the potential for evolutionary responses to environmental change. This challenge is more difficult to overcome, since field estimates of genetic variation within and among populations are clearly not feasible for all taxa. Nevertheless, the current study highlights how these assessments can be undertaken using model organisms such as *Drosophila*.

## ACKNOWLEDGEMENTS

We are very grateful to Peter Alexander, Chris Clinton, Ciara Mann, Amanda McGeady and Lara Meade for technical assistance. Thank you to Roger Butlin for helpful discussions on experimental design, and to James Buckley and three anonymous reviewers, whose insightful comments on earlier versions have greatly improved this manuscript. This work was funded by a Natural Environment Research Council Standard Grant to JRB. The authors declare no conflict of interest.

## Supporting Information captions

**Table S1.** Location and environmental variation of altitudinal gradients where *D. birchii* was collected between 2010 – 12, including altitudinal range, total length (the straight-line distance between the top and bottom of each gradient in km), number of sites sampled, ranges of environmental variables (Mean daily temperature (MDT); Mean daily minimum temperature (MDT_min_); Mean daily maximum temperature (MDT_max_); Mean daily temperature difference (MDT_diff_); Mean daily humidity (MDH)), *D. birchii* density, density of other species from the *serrata* species complex (non-*birchii* density), and productivity in cages (only assessed in 2012). For each environmental variable, density and cage productivity, the range shown is the mean at the lowest altitude site to the mean at the highest altitude site. Density of *D. birchii* and other *serrata*-complex species were not estimated in 2012. The difference in mean temperature between the most northerly gradient (Mt Lewis) and the most southerly gradient (Paluma) was less than the temperature difference seen within most of the altitudinal gradients.

**Table S2:** Linear regressions of each environmental variable measured during 2010–2012 on (a) altitude for each gradient, and (b) altitude, latitude and their interaction across the entire sampled range. Shown is the slope, with the Standard Error (SE) in brackets, of the regression line between each environmental variable and altitude/latitude, and the *R^2^* value indicating the proportion of variation explained by the model. *N* is the number of sites sampled. Symbols indicate the significance of each factor in the model: ^***^ *P* < 0.001, ^**^ 0.001 ≤ *P* < 0.01, ^*^ 0.01 ≤ *P* < 0.05, † 0.05 ≤ *P* < 0.1, ^NS^ *P* ≥ 0.1. Significant associations are highlighted in *italics*.

**Table S3:** Correlations between environmental variables included as predictors of *D. birchii* field abundance (below diagonal) and *p*-values indicating significance of correlations (above diagonal). All correlations were highly significant, even at a very conservative Bonferroni-corrected significance threshold of *P* = 0.003.

**Table S4.** Loadings of each environmental variable measured along the four gradients on the first two Principal Components (PCs) from a Principal Component Analysis. The first two PCs together accounted for 89 % of the variation in these variables.

**Table S5.** Variation in productivity among isofemale lines (nested in source population) from Mt Edith and Paluma when reared in the laboratory. Productivity variation was analysed using Generalised Linear Mixed Models (GLMMs), run in the *R* package *glmmADMB*, specifying zero-inflation, and a negative binomial distribution with a log link function. Source population (indicating which of the four populations within a gradient the line came from) was included as a fixed factor and maternal isofemale line (‘Line’), nested within source population, was included as a random factor. Significant effects are denoted in *italics*. The significance of fixed effects was evaluated using a χ^2^ test, and of random effects using a likelihood-ratio test comparing models with and without the term included. Separate analyses were conducted for the two gradients. Productivity was measured as the mean number of offspring per female produced from controlled crosses in the laboratory. Sites at both gradients differed significantly in their productivity in the lab (but not in the field; see Table 2). Estimates of among-line variance in productivity at both gradients were much higher in the laboratory than in the field (*cf* Table 2), and this variance was significant at Mt Edith.

**Table S6.** Results of linear models to test how well mean fitness in cages (cage productivity) predicts local abundance in the field. Separate analyses were performed for Mt Edith and Paluma, the two gradients where caged transplants were performed. All fitness and abundance data were standardized to mean = 0; standard deviation = 1 prior to analysis. Shown are the slopes of the regressions of cage productivity on local abundance at each gradient, with the standard error of this estimate in brackets, the *t*-value for the analysis, and the significance of each test.

**Figure S1.** Laboratory crossing scheme to generate lines used in cage transplants from each of the four source populations from each gradient. Females from each of the (up to five) isofemale lines from each population were mated with males from each of the other lines from the same population, excluding crosses between flies from the same line. Each line cross combination was replicated three times. Offspring of females from the same isofemale line were then combined and used in the cage transplant experiment. There was substantial variation in the number of offspring generated by line combinations (see Table S4; Figure S3), therefore the number of lines available for transplant varied among populations and sites. Exact numbers of lines transplanted to each site are shown in the table within Figure S3.

**Figure S2.** Schematic illustrating design of caged transplant experiment. Bold lines show transplants among the sites of origin: solid lines indicate transplants back into each population’s home site; large dashed lines indicate transplants to the other site of origin at the same end of the gradient; and small dashed lines indicate transplants to the sites of origin at the opposite end of the gradient. Dotted lines indicate transplants to intermediate sites along the gradient (I-1 – I-5). Transplants originating from low altitude sites (LOW 1 and LOW 2) are in red, and from high altitude sites (HIGH 1 and HIGH 2) are in blue. To improve clarity, only one set of arrows depicts transplants from each end of the gradient to sites along the gradient, but lines from both populations of origin were transplanted in each case. At Paluma, only high altitude lines were transplanted to intermediate sites, but all lines were transplanted to sites at gradient ends. *The number of lines transplanted to each site varied because some lines did not produce sufficient offspring (see Methods). The table shows the exact number of lines and cages transplanted into each site at each of the two gradients.

**Figure S3.** Comparison of temperatures measured inside field cages using iButtons (filled symbols) and outside cages at field sites using Tinytag Data Loggers (open symbols) along the two gradients where field transplant experiments were undertaken: Mt Edith (left) and Paluma (right). There was no significant difference between the estimates of mean daily temperature (MDT) (squares; *t* = −1.50, *P* = 0.142), or mean daily maximum temperature (MDT_max_)(circles; *t* = 0.367, *P* = 0.716) inside and outside cages, although mean daily minimum temperatures (MDT_min_) measured using iButtons inside cages were lower than those measured outside cages using Data Loggers (triangles; *t* = − 5.78, *P* = 1.37 × 10^-6^). There was no significant difference between iButtons and Data Loggers in the change in temperature with respect to altitude for any of the three measures: MDT (*t* = 0.76, *P* = 0.452), MDT_min_(*t* = 1.24, *P* = 0.222) or MDT_max_(*t* = 0.86, *P* = 0.396).

**Figure S4.** Mean productivity of each of the four source populations from Mt Edith (left) and Paluma (right) in laboratory crosses. Error bars are standard errors across mean productivities of the (up to five) lines within each source population. At one of the high altitude sites from Mt Edith (High1), only one of the five lines produced offspring in laboratory crosses. Different letters indicate significantly different productivity means of populations within a gradient.

**Figure S5.** Mean productivity (estimated as the mean number of offspring per female) of each of the four source populations from Mt Edith (left) and Paluma (right) in cages transplanted to sites along altitudinal gradients. Productivities are averages across cages within and among sites for each source population. Error bars are standard errors across mean productivities of the (up to five) lines within each source population. At one of the high altitude sites from Mt Edith (High1), only one of the five lines produced offspring in laboratory crosses, therefore only a single line could be transplanted from this site. There was no significant difference among source populations at either gradient (see Table 2).

